# Equine intestinal O-seroconverting temperate coliphage Hf4s: genomic and biological characterization

**DOI:** 10.1101/2021.06.09.447823

**Authors:** Eugene E. Kulikov, Alla K. Golomidova, Alexandr D. Efimov, Ilya S. Belalov, Maria A. Letarova, Evelina L. Zdorovenko, Yuriy A. Knirel, Andrey S. Dmitrenok, Andrey V. Letarov

## Abstract

Tailed bacteriophages constitute the bulk of the intestinal viromes of the vertebrate animals. However, the relationships between lytic and lysogenic lifestyles of the phages in these ecosystems are not always clear and may vary between the species or even between the individuals. The human intestinal (fecal) viromes are believed to be dominated by temperate phages, while in the horse feces the virulent phages are more prevalent. Almost all the isolates of horse fecal coliphages are virulent. Phage Hf4s is the first temperate equine intestinal coliphage characterized. It was isolated from the horse feces on the indigenous equine *E. coli* 4s strain. It is a podovirus, related to *Lederbergvirus* genus (including the well–characterized *Salmonella* phage P22). Hf4s recognizes the host O antigen as its primary receptor and possesses a functional O-antigen seroconversion cluster that renders the lysogens protected from the superinfection by the same phage and also abolishes the adsorption of some indigenous equine virulent coliphages, such as DT57C, while the other phages, such as G7C or phiKT retain the ability to infect *E. coli* 4s (Hf4s) lysogens.

**Importance:** The relationships between virulent and temperate bacteriophages and their impact on high-density symbiotic microbial ecosystems of the animal are not always clear and may vary between the species or even between the individuals. The horse intestinal virome is dominated by the virulent phages, and Hf4s is the first temperate equine intestinal coliphage characterized. It recognizes the host O antigen as its primary receptor and possesses a functional O-antigen seroconversion cluster that renders the lysogens protected from the superinfection by some indigenous equine virulent coliphages, such as DT57C, while the other phages, such as G7C or phiKT retain the ability to infect E. coli 4s (Hf4s) lysogens. microbial viruses in the mammal intestinal ecosystems.

## 1. Introduction

Bacteriophages are now widely recognized as an essential component of most of the natural microbial systems including the symbiotic microbial communities of the human or animal body (1–3). Nowadays most of the work on intestinal bacteriophages is performed using holistic approaches such as metagenomic sequencing of the viral DNA or RNA. However, the data generated by the -omics approaches are currently hard to interpretation even in humans for which the diversity of intestinal bacteriophages has been extensively investigated over the last two decades (4, 5). In other mammal species, the vast majority of the viral genomic sequences represent so-called “genomic dark matter” matching no close relatives in databases of sequenced bacteriophage genomes (6). Therefore, the culture-based research is still relevant for better understanding of the bacteriophage ecology in the gut (1, 5). Although substantial advancements were made in culturing phages of the major microbial groups of the human intestinal microbiome including highly represented crAssphage-like viruses (7, 8), *E. coli* remains a valuable model organism. The study on the diversity and dynamics of the indigenous intestinal coliphages provides useful opportunities to and provide insights into the mechanisms of the ecological interactions and the modes of the evolution of the intestinal bacteriophages and their hosts (9–11).

The most prolific bacteriophage group comprising the vast majority of the known viral isolates is the tailed phages (order Caudovirales). Among these are virulent and temperate bacteriophage species. The virulent bacteriophages immediately replicate in any cell they infect, consequently killing all the successfully infected cells. In contrast, the temperate phages take lytic or lysogenic decision upon every cell infection. A fraction (in most cases – the majority) of the host cell undergoes immediate lysis, while the rest of them are converted into the lysogenic clones in which the bacteriophage genome gets integrated into the bacterial chromosome and exists in the form of prophage (12). In some phages the prophage exists as a circular (e.g. in phage P1 (13)) or linear (phage N15(14)) episome. The prophage-harboring cells referred to as lysogenic may undergo prophage induction that leads to the prophage excision and lytic development. However, the lysogenic cells are normally immune to the superinfection by the same phage due to the presence of the repressor(s) blocking the expression of the newly injected viral genome. Some prophages also encode proteins modifying the lysogenic cell surface to prevent adsorption or entry of the new particles of the same phage and also of some other phages (15), including the modifications of the O antigen leading to serotype conversion (16). Besides the cell surface alterations, the temperate phages may encode other conversion genes that are expressed in the lysogens providing the cell with some novel features increasing the fitness under particular conditions (12, 17). The prophages may also participate in the cellular gene expression control circuits (18) and contribute to bacterial fitness by some other yet poorly understood mechanisms (19, 20). Therefore, the prevalence and the nature of the temperate phages in particular natural viral communities (viromes) may have diverse and profound implications for the life of the corresponding microbial biota.

Human fecal viromes are believed to be dominated by the temperate phages (21). It was demonstrated that the pool of the cultured *Escherichia coli* phages found in the stool of healthy humans is also mainly represented by the temperate viruses ((9, 22), see also (1)), although the fraction of the virulent phages increased in the patients with diarrhea (22, 23). The available data on the abundance of the virulent and temperate phages in the feces of other mammal species is scarce. However, in the metagenomes of the horse feces the fraction of the contigs attributed to temperate phages was moderate (6). The coliphages isolated from the pool of the free phage particles of horse feces were almost entirely represented by the virulent phages ((11, 24–28) and our unpublished observations). Up to our knowledge, the isolation of the temperate coliphages from the horse feces was never reported in the literature.

Here we describe the genome and biological characterization of Hf4s – the first temperate *E. coli* phage obtained from the horse feces. Hf4s is an O antigen dependent virus infecting an indigenous equine E. coli strain 4s. The lysogens harboring Hf4s undergo the serotype conversion due to additional lateral glycosylation of O antigen (O-unit) that leads to the Hf4s adsorption exclusion and also provides resistance to some of the virulent bacteriophages infective for *E. coli 4s.*

## 2 Materials and methods

### 2.1 Bacterial and bacteriophage strains

All the strains used were from our laboratory collection. *E. coli* 4s strain producing О22-like O antigen was previously obtained from the horse feces sampled at the same location and characterized by us (29). *E. coli* 4s mutants were described in (29, 30). These were: 4s GTR, lacking the lateral glucose residue on the O-unit, 4sI (*wcl*K^−^), lacking the O antigen O-acetylation and 4sR (*wcl*H^−^) defective for O-unit assembly and, therefore, completely devoid of O antigen.

Bacteriophage G7C is an N4-related podovirus, isolated from the horse feces (24). This phage is strictly specific to *E. coli* 4s strain depending on the O-acetylated O antigen for the host cell recognition (31). Clones selected for resistance to this virus are lacking O antigen synthesis or O antigen O acetylation (29, 30).

Phage phiKT is also a horse-derived podovirus distantly related to the phage T7. It strictly depends on O antigen for infection, although it is not sensitive for the O-acetylation (30).

Phage DT57C is closely related to the phage T5 (26). It recognizes BtuB protein as its secondary receptor, but the O antigen of *E. coli* 4s prevents it from the direct interaction with the protein receptor. To infect the wild type *E. coli* 4s, this phage recognizes O-polysaccharide by its lateral tail fiber protein LtfA (32). Phage FimX (DT571/2-ltfA^−^) is a mutant of the bacteriophage DT571/2, closely related to DT57C. This mutant lacks the lateral tail fibers and infects only rough mutants of *E. coli* 4s and the mutants carrying non-acetylated O antigen (the lack of the O-acetylation leads to the decreased O antigen production; (30)) This features allows to use the phage FimX as a probe to assess the nonspecific protection of the bacterial cell surface by the O antigen.

### 2.2 Bacteriophage isolation and cultivation

Phage Hf4s was isolated from the sample of the horse feces collected in the stable of equestrian center in Neskuchny sad in Moscow, Russia in 2015, following the procedure described in (11). The bacterial host used for the phage isolation was *E. coli* 4s. The bacteriophage was purified by repeated single plaque isolation. To grow high-titer phage stock, 25 ml liquid culture of *E. coli* 4s host was inoculated by a single plaque and grown at 37° C with vigorous agitation overnight.

### 2.3 Phage morphological examination

The bacteriophage particles morphology was revealed by transmission electron microscopy (TEM) of the particles negatively stained with 1% uranyl acetate in methanol. The microscopy was performed as described earlier (32).

### 2.4 Genomic sequencing

The viral DNA was extracted from the DNAse/RNAse-treated lysate as described in (25) and submitted for genomic sequencing. The sequencing was using an IonProton sequencer system (Applied Biosystems, USA) with 45-fold coverage and a median length of 201 bp. The raw reads from the run were then combined and filtered using the spectral alignment error correction tool SAET 3 (Applied Biosciences, USA). The genome sequence was assembled using SPAdes (33). The genome was then annotated manually using CLC-genomics 6.0 software package.

### 2.5 Bioinformatic analysis

The bacteriophage genomes related to the bacteriophage Hf4s were retrieved from GenBank using tblastn (34). Briefly, concatenated protein sequences corresponding to three distinct genome regions (0-16042, 16043-29499, and 29500-39390) were used as query sequences in three separate blast runs. All Hf4s concatenated proteins were used as a query in a separate blast run. Among more than 200 top hits from all blast runs 25 sequences providing the query coverage greater than 35% were selected for further analysis. In the selected sequences prophages were identified with Phigaro (35). All selected nucleic sequences turned out to be prophages in bacterial genomes. The same approach was used for selecting viral genomes, but this time only among the collection of Caudovirales (taxid: 10239) sequences. The sequences in which the regions aligned with the query covered more than 35% of the later were selected for further analysis. Sequences of prophage sequences and phage genomes related to Hf4s genome described in Tables S1 and S2 respectively. Identified prophage and free viral genome sequences were visualized with Easyfig (36) with tblastx alignment.

Selected prophage sequences were aligned with Mauve (37). Since only 10 protein-coding genes were shared among all 25 viruses (none of them in the region corresponding to 11514-34507 bp of Hf4S genome), we didn't investigate details of phylogeny for individual genes.

All pairwise comparisons of all the amino acid sequences encoded by free phages and prophages were conducted using the Genome-BLAST Distance Phylogeny (GBDP) method (38) under settings recommended for prokaryotic viruses (39). We used T4 phage genome as outgroup, since several siphoviruses were related to Hf4s with blast search along with podoviruses.

The resulting intergenomic distances were used to infer a balanced minimum evolution tree with branch support via FASTME including SPR postprocessing (40) using D6 formula (39), since it produces better supported tree. Branch support was inferred from 100 pseudo-bootstrap replicates each. Trees were rooted at the midpoint (41) and visualized with FigTree (42).

Taxon boundaries at the species, genus and family level were estimated with the OPTSIL program (43), the recommended clustering thresholds (39) and an F value (fraction of links required for cluster fusion) of 0.5 (44).

### 2.6 Lypopolysacharides (LPS) profiling

SDS-PAGE profiling of the LPS of bacterial strains was performed as it described in (30).

### 2.7 Bacteriophage adsorption curve determination

The adsorption curve was determined as described in (45). Briefly: the early-log phase (OD_600_ = 0,2) culture of *E. coli* 4s was grown in LB medium at 37 °C with agitation. Appropriately diluted phage stock phage was added to 1 ml of the culture in the 1,5 ml Eppendorf tube up to the final of about 10^5^ PFU ml^−1^. The mixture was incubated at 37°C in the thermoshaker. At the defined time points the samples of 10 mkl were taken and immediately diluted in 990 mkl of the physiological saline to slow down the page adsorption. The diluted samples were centrifuged in a tabletop microcentrifuge at 12 000 g for 1 min. The 50 mkl aliquots of the supernatant were plated out to count the PFU numbers. The whole experiment was triplicated. The adsorption constant was calculated between the points 1 min and 3 min.

### 2.7 Isolation of the O-polysaccharide and NMR spectroscopy

The LPS was extracted from *E. coli* cells using phenol-water extraction protocol (46).

The O-polysaccharide was obtained by hydrolysis of the lipopolysaccharide with aqueous 2% HOAc for 2 h at 100 °C, a lipid precipitate was removed by centrifugation (13,000*g*, 20 min), and the carbohydrate portion was fractionated by gel-permeation chromatography on a column (56 × 2.6 cm) of Sephadex G-50 Superfine (Healthcare) in 0.05 M pyridinium acetate buffer pH 4.5 monitored with a differential refractometer (Knauer, Germany).

A polysaccharide sample was deuterium-exchanged by freeze-drying twice from 99.5% D_2_O and then examined as a solution in 99.95% D_2_O. ^1^H- and ^13^C NMR spectra were recorded using a Bruker Avance II 600 MHz spectrometer (Germany) at 50 °C using internal sodium 3-trimethylsilylpropanoate-2,2,3,3-d_4_ (δ_H_ 0.0, δ_C_ –1.6) as reference. Two-dimensional NMR experiments were performed using standard Bruker software, and Bruker TopSpin 2.1 program was used to acquire and process the NMR data. A spin-lock time of 60 ms and a mixing time of 100 ms were used in TOCSY and ROESY experiments, respectively. An HMBC experiment was run with a 60-ms delay for evolution of long-range couplings.

### 2.8 Accession number

The whole genome sequence of the bacteriophage Hf4s was deposited to GenBank under the accession number MT833387.

## 3 Results

### 3.1 Bacteriophage Hf4s isolation and general characterization

Bacteriophage Hf4s was isolated in 2015 from the sample of the horse feces collected at the Equestrian center in Neskuchny sad (Moscow, Russia) using *E. coli* 4s strain as the isolation host. Noteworthy, *E. coli* 4s was originally isolated from the same location in 2006 (11). The plaques of Hf4s phage had unusual morphology, being larger than most of the coliphages plaques usually observed upon the plating of the extract of horse feces, appearing as multiple concentric turbid rings spaced by clear zones. The multiple-rings morphology could be better seen around the drop of the diluted phage suspension applied to the lawn (eye-like plaque morphology, Fig. 1A). If a drop of the phage suspension was applied on bacterial lawn, the turbid growth inhibition zone was formed, that was surrounded by 2-3 concentric rings similar to the rings observed around the isolated plaques (Fig. 1A). This plaque morphology was suggestive for the lysogenic lifestyle of the phage. To prove that the bacterial growth inside the plaque or lysis spot is due to formation of the lysogens, we re-suspended the material taken from the middle of the inhibition zone in the viricidal solution (47) to remove all the external phage and plated the serial dilutions onto LB plates to obtain the individual subclones. Twenty subclones were transferred with the toothpicks onto fresh LB plate and onto the plate with the soft agar inoculated by the *E. coli* 4s strain. All the clones grew well on LB plate and all produced the zone of the inhibition of the lawn growth around the colonies of the plate with *E. coli* 4s due to phage production. The ability to produced phage maintain through three sequential passages with viricidal treatment. Therefore, we concluded that Hf4s is a temperate phage and the obtained clones are its lysogens designated as *E. coli* 4s (Hf4s).

**Figure 1.**
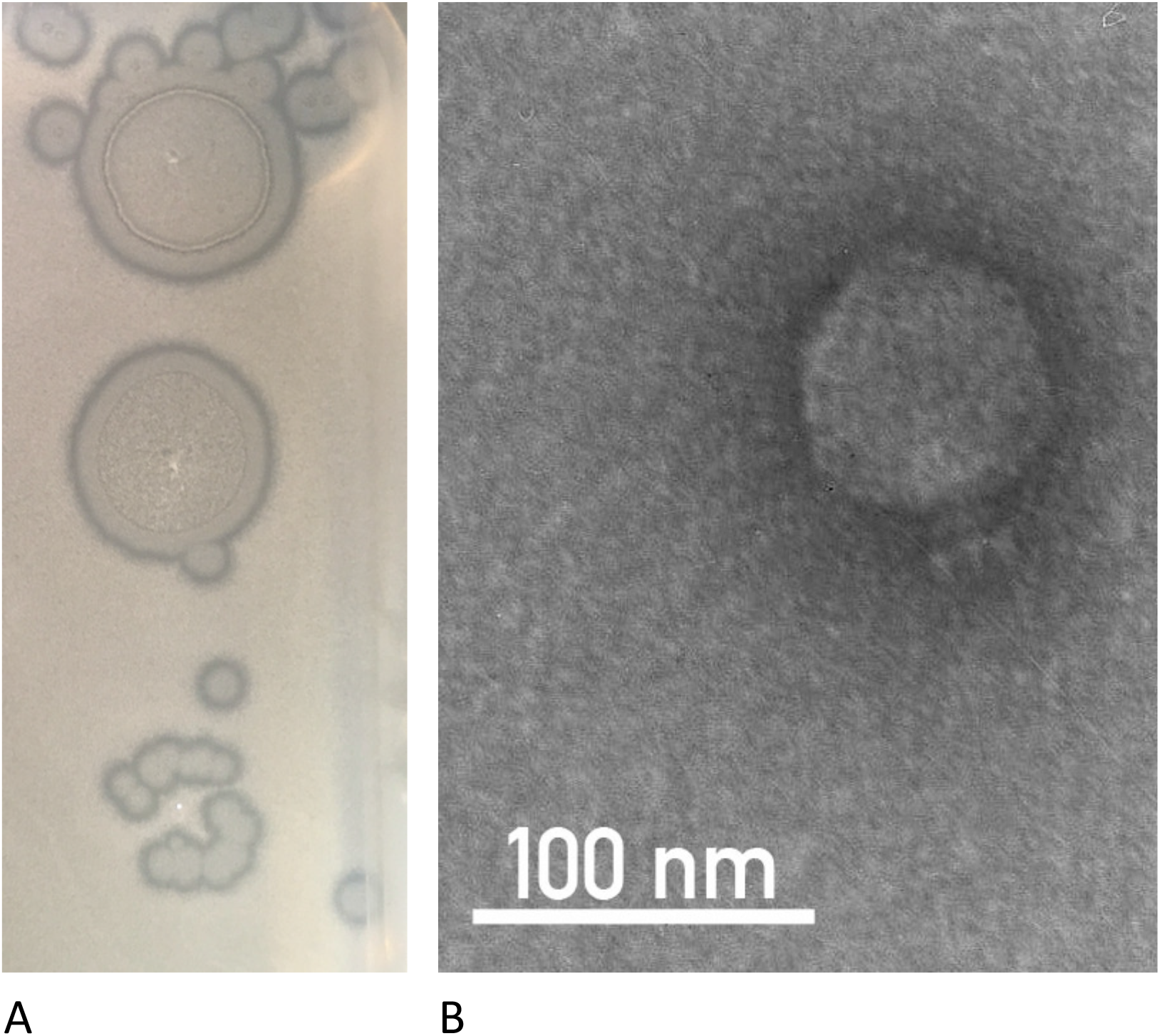
A – morphology of the bacteriophage Hf4s plaques on the lawn of *E. coli* 4s host strain. The concentric rings are better visible around the spots of concentrated phage suspension. B – the Hf4s virion morphology.

The infection of the liquid culture by an isolated Hf4s plaque led to rapid (typically within 4-5 h) lysis of the culture with flocculation, yielding high titer of the bacteriophage (typically about 10^11^ PFU ml^−1^). However, the overgrowth of phage-resistant bacteria (apparently lysogens) was observed almost immediately. The titer of the lysate remains high even if it is incubated overnight, despite the massive overgrowth of the lysogens.

To reveal the phage morphology, we performed TEM examination of the negatively stained virus. Phage Hf4s was found to be a small podovirus with an isometric icosahedral head of 65 nm in diameter and a short tail with the tail spikes (Fig 1B).

### 3.2 Phage Hf4s genome

Whole genome sequencing of the Hf4s bacteriophage yielded a single contig of 39 390 b.p. length. The ORF prediction using GenMark software resulted in 61 potential ORFs in the Hf4s genome (Fig. 2). The annotation was performed manually using blastP and HHpred to predict the function of the ORFs. The overall genome structure was typical to P22-related lambdoid phages. Left genome is occupied by the morphogenetic gene cluster (presumably an operon), while the lysogeny control and lysogenic conversion modules are located in the middle followed by the late genes, such as antiterminator protein Q gene and the host cell lysis module.

**Figure 2.**
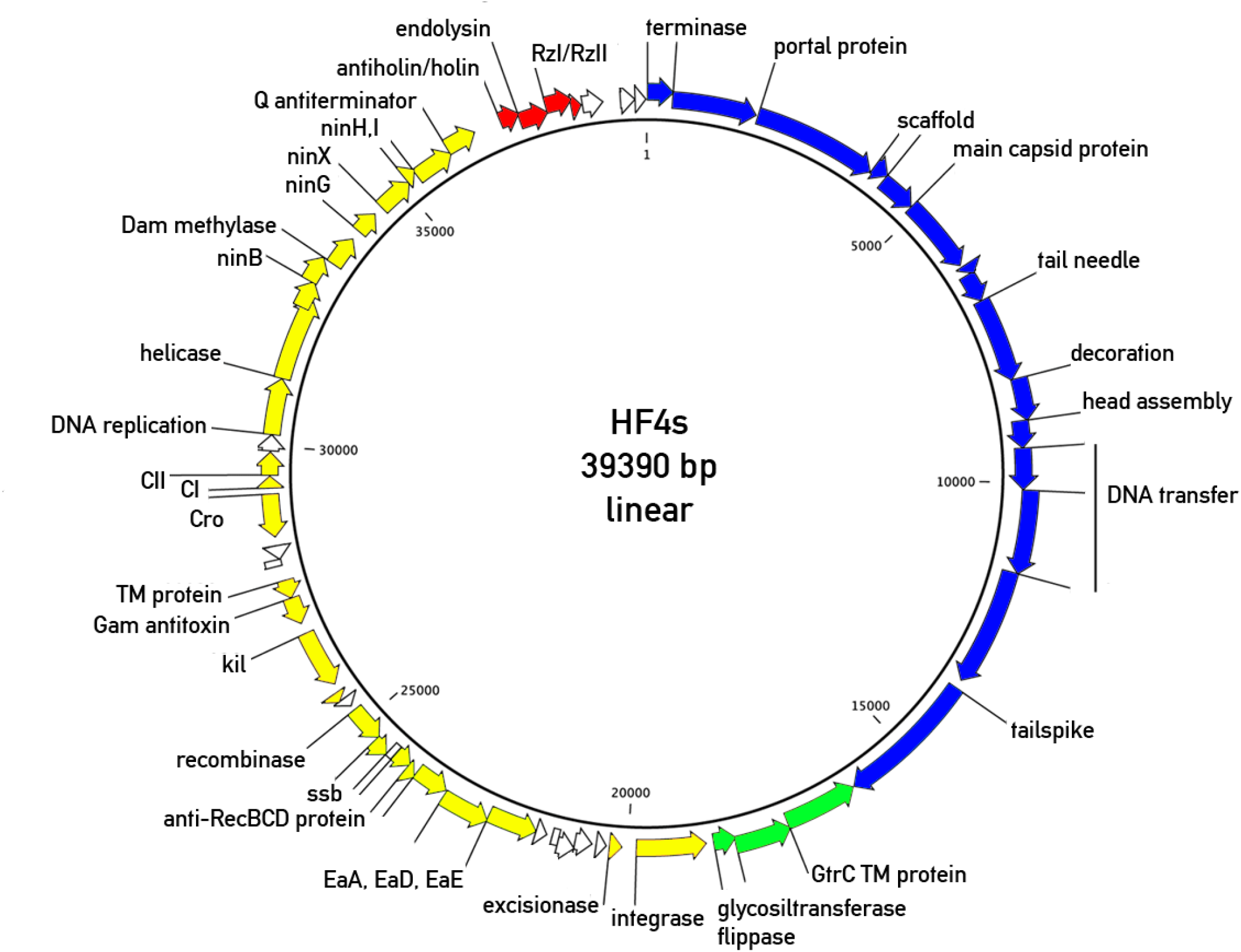
Genomic map of the bacteriophage Hf4s. Yellow – middle and early gene cluster, lysogeny control and DNA replication genes. Red, white and blue – late cluster, including the host lysis genes (red), genes of unknown function (white) and morphogenetic genes (blue). Green – O-seroconversion cluster.

The NCBI blastn search against the database of the viral sequences showed that Hf4s contained genomic fragments closely related to the sequences of known temperate bacteriophages. The best hits were for the lambdoid podoviruses HK60, Sf6 and N7, distantly related to the well-characterized *Salmonella* phage P22, all these phages belonging to the *Lederbergvirus* genus (Fig. S1). The level of the nucleotide identity of the aligned sequences was very high, reaching 97%. However, both in case of prophages or viral genomes large fraction of the Hf4s genomic sequence was not related to the retrieved sequences indicating the mosaic organization of the genome that is typical for the lambdoid phages (48). The left 8.5 kbp fragment of the Sf6 genome was aligned to the Hf4s sequence with 97% nucleotide identity. Since this fragment contains the genes for the small and large terminase subunits that are responsible for the *pac*-site recognition and cleavage, we set the left genomic end at the same site as in the phage Sf6 genome. The RFLP analysis of HF4s genomic DNA using BamHI REase revealed the presence of ~6.700 bp long DNA fragment in substoichiometric quantity vs. rest of fragments (data not shown). This implies that HF4s phage uses pac-site packaging strategy yielding limited circular permutations. The length of this fragment is consistent with the presumable left-end position of the genome.

To better elucidate the genetic relationships of Hf4s with other viruses, additional nucleotide sequences related to the Hf4s genome were retrieved from the total nucleotide (nr) database using tblastn algorithm with the concatenated phage proteins sequence used as a query (see Material and methods for detail). The hits were to multiple prophages from different Enterobacteriaceae genomes. Interestingly, the closest matches were for *Salmonella enterica* prophages though several *E. coli* prophages were also found to share homologous regions with Hf4s (Fig. 3, 4 and Tables S2 and S3). Most of the prophages identified had the size close to the genome size of Hf4s (~40 kbp) and belong to podoviruses. However three prophages of much greater size were identified, comprising two 62 - 63 kbp long (Fig. 4) and one 112 kbp prophage. These prophages also belong to the podoviruses sharing with Hf4s capsid genes and some of the replication, transcription control and lysogeny genes (Fig. 4.). It has to be mentioned that the boundaries of all the prophages including the bigger ones were determined from Phigaro software output. The manual inspection of these sequences regions of the large prophages seem to contain large number of potentially bacterial genes that may be suggestive for either wrong identification of the prophages ends or for ongoing process of the degradation of defective prophages washing the boundaries with the host genetic material.

**Figure 3.**
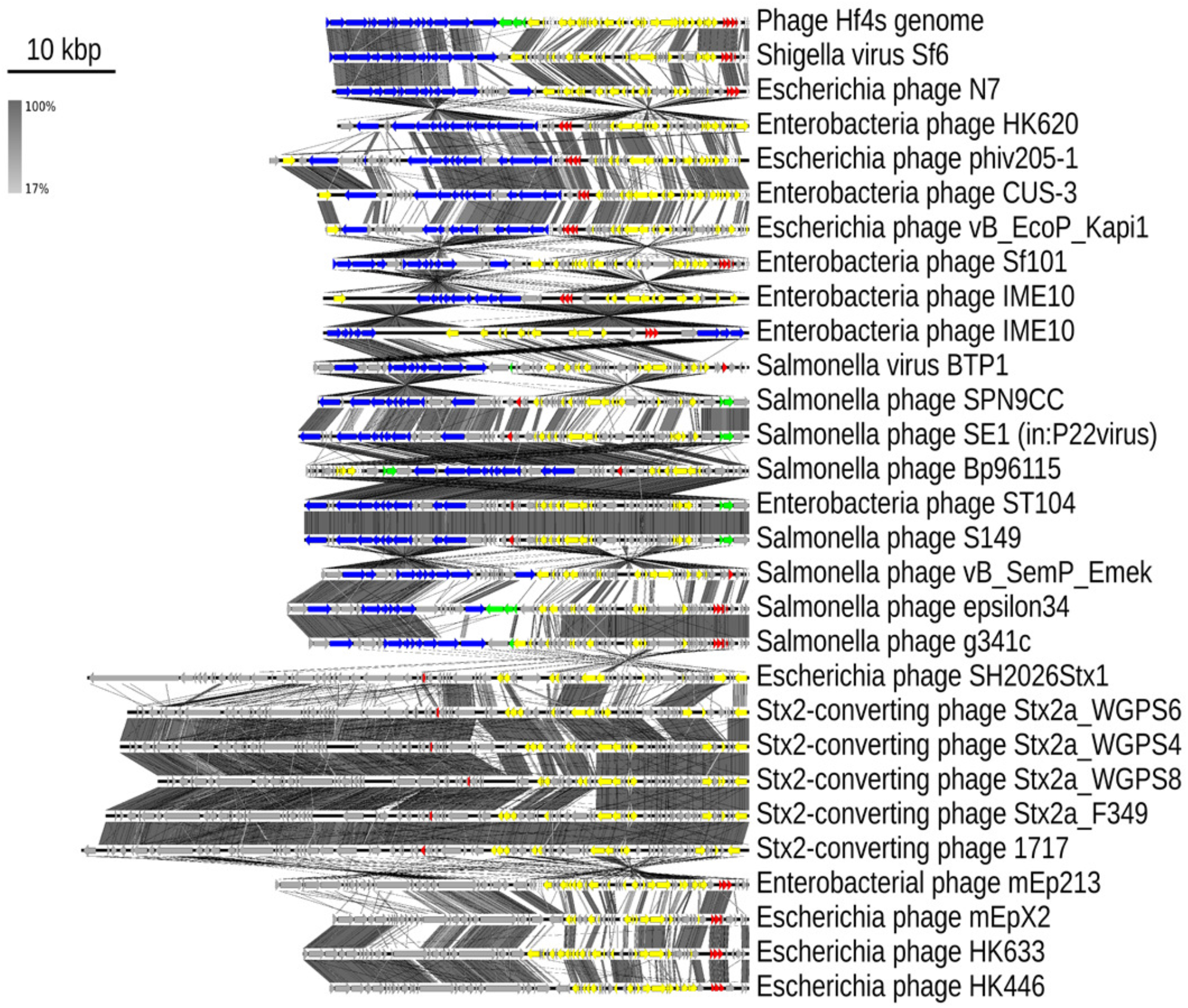
Comparison of the bacteriophage genomes related to Hf4s. The color code for the genes showing the similarity to the phage Hf4s genes is the same as on Fig. 2. The unrelated ORFa are shown in grey.

**Figure 4.**
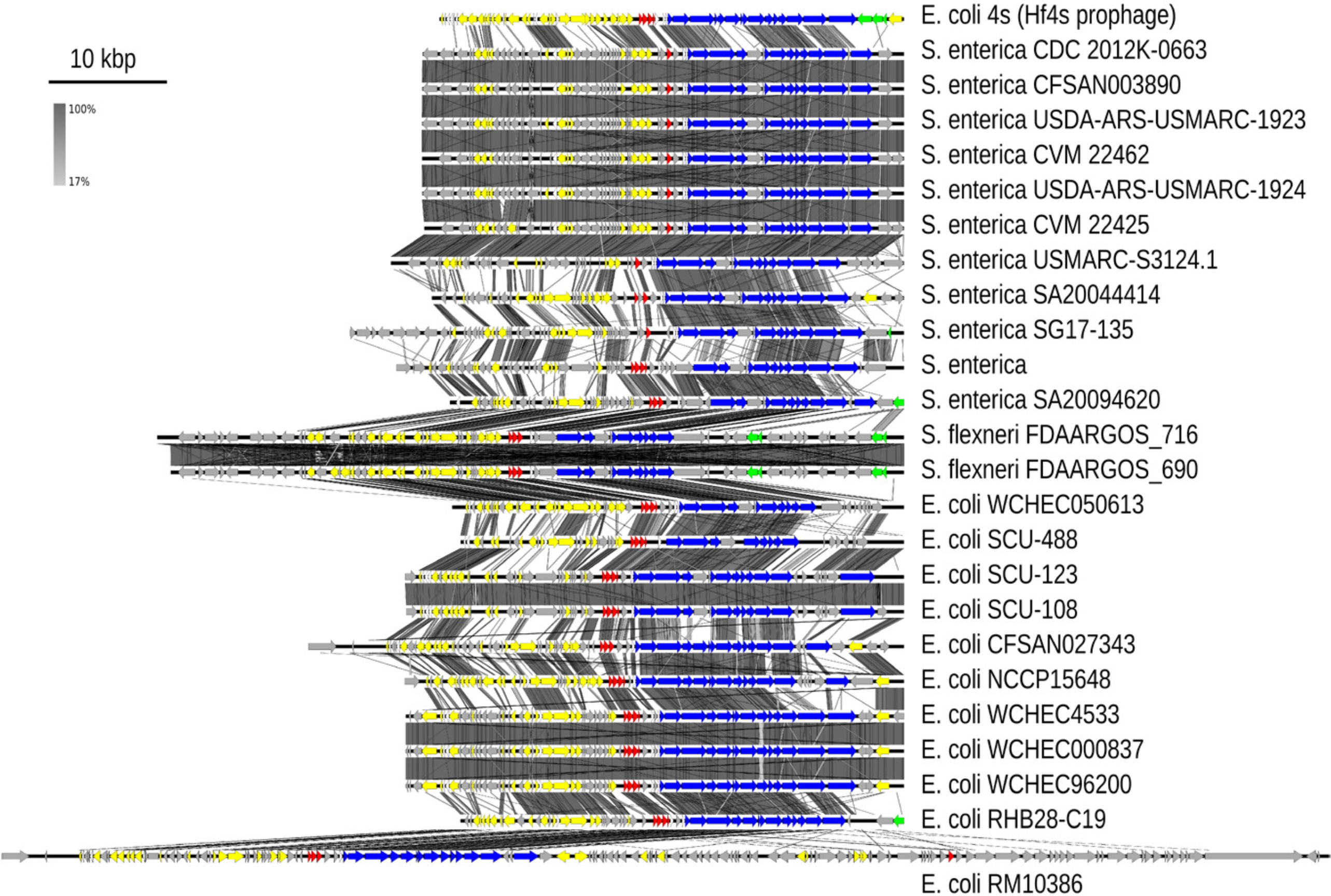
Comparison of the prophage sequences related to Hf4s. The genome sequence of Hf4s was *in silico* circularized and re-opened at the predicted att site (similar to the *att* site of the closest related prophage from the *Salmonella enterica* CDC 2012K-0663 strain) The color code for the genes showing the similarity to the phage Hf4s genes is the same as on Fig. 2. The unrelated ORFa are shown in grey.

The phylogenetic GBDP trees were inferred using FASTME software with the D6 formula. The average support value of 52 % was obtained (Fig 5). The numbers above branches are GBDP pseudo-bootstrap support values from 100 replications. The branch lengths of the resulting VICTOR trees are scaled in terms of the D6 distance formula (39).

**Figure 5.**
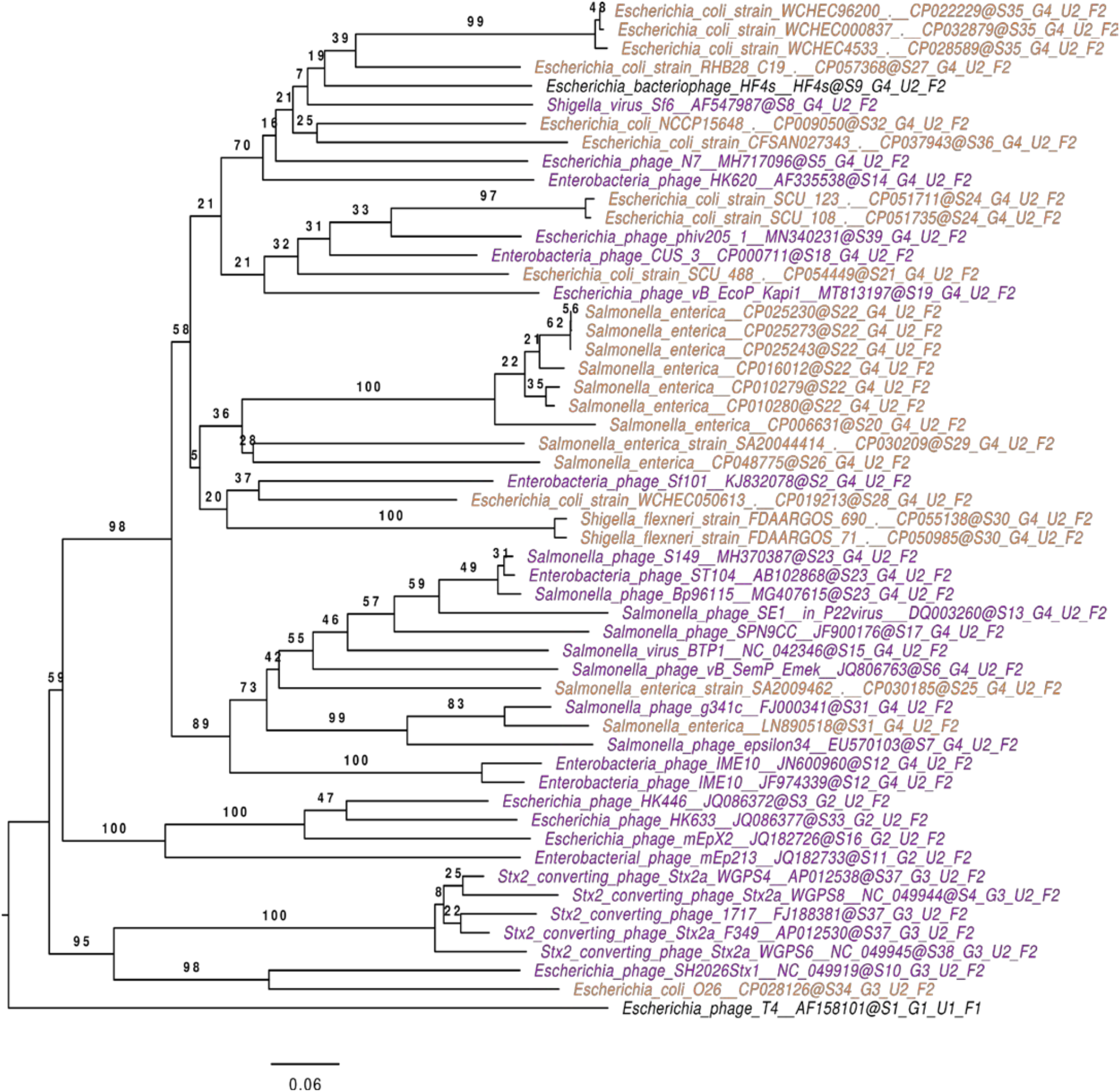
Phylogeny of Hf4s – related prophages and phages as revealed by VICTOR algorithm. Phage T4 used as an out-group.

The OPTSIL clustering yielded 39 species clusters. At the genus level, four clusters were inferred. Number of families and subfamilies were two, separating out-group phage T4. Therefore we concluded that 42 out of 53 sequences related to Hf4s could be classified in a single genus-level, roughly corresponding to the established *Lederbergvirus* genus.

### 3.4 Host cell recognition by Hf4s bacteriophage

In our previous work (30) we tested Hf4s (among a number of other *E. coli* 4s viruses) for infectivity against the series of the mutants of *E. coli* 4s strain described by us earlier6 deficient of different stages of O antigen or core oligosaccharide synthesis. We found that the phage efficiently forms the plaques on the strain lacking the lateral glucose modification (GTR), the growth was significantly inhibited on the strain lacking the O-acetylation of the O units (4sI), and the strains lacking the O antigen backbone were completely resistant to the Hf4s infection (Table 1). Therefore, we concluded that *E. coli* 4s O antigen is a primary receptor of Hf4s and the interaction with this receptor is essential for subsequent steps in the infection (i.e. the infection *via* the direct recognition of the secondary receptor is not possible).

**Table 1.**
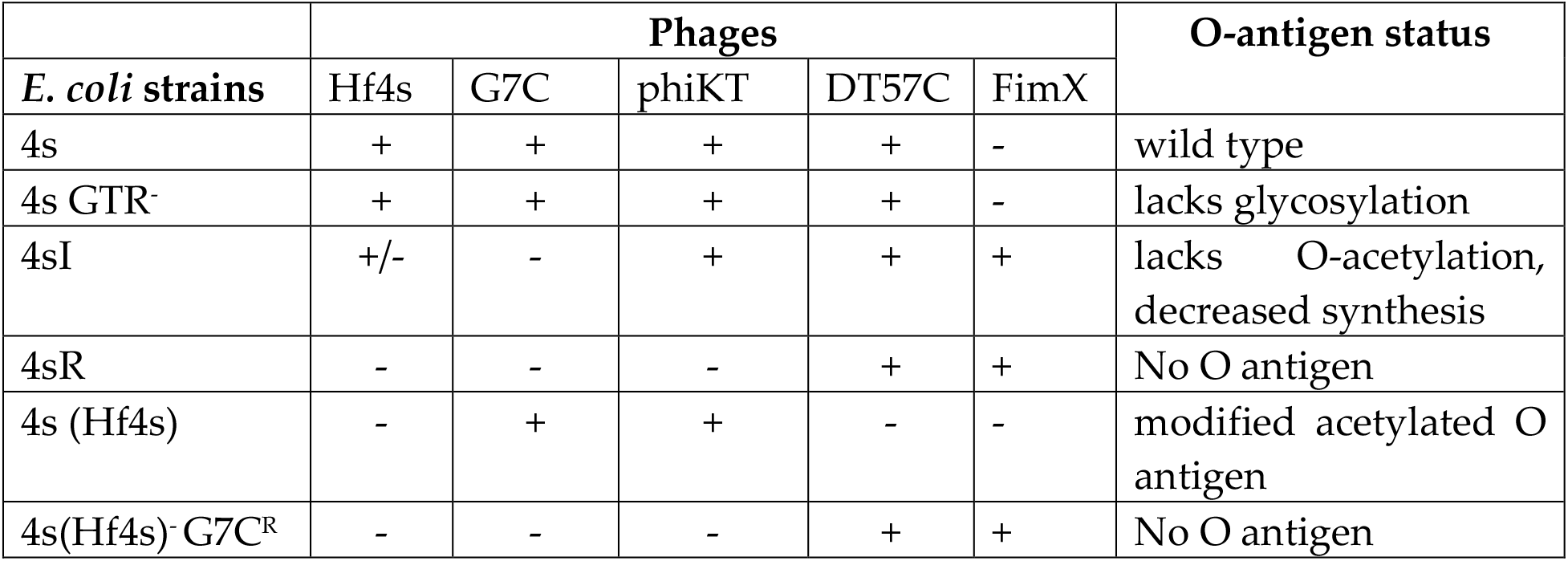
Sensitivity of *E. coli* 4s derivative strains to the bacteriophages. “+”– efficient plaque formation, “-“ – no visible growth, “+/−“ – no plaques formed, but the lawn inhibition spot is formed if a drop of the concentrated (10^8^ PFU ml^−1^) phage suspension is applied. The O antigen status of E. coli 4s and its mutants were determined previously (30), the O antigen of the Hf4s lysogens was determined in this work.

We determined the curve of the bacteriophage Hf4s adsorption on *E. coli* 4s cells (Fig 6). The adsorption constant was estimated from the data of this experiment as 6×10^−9^ ml min^−1^. High adsorption constant value, close to the theoretically predicted limit of about 10^−8^ ml min^−1^ (45) is in good agreement with the identification of the O antigen, a highly represented molecule, as the Hf4s primary receptor.

**Figure 6.**
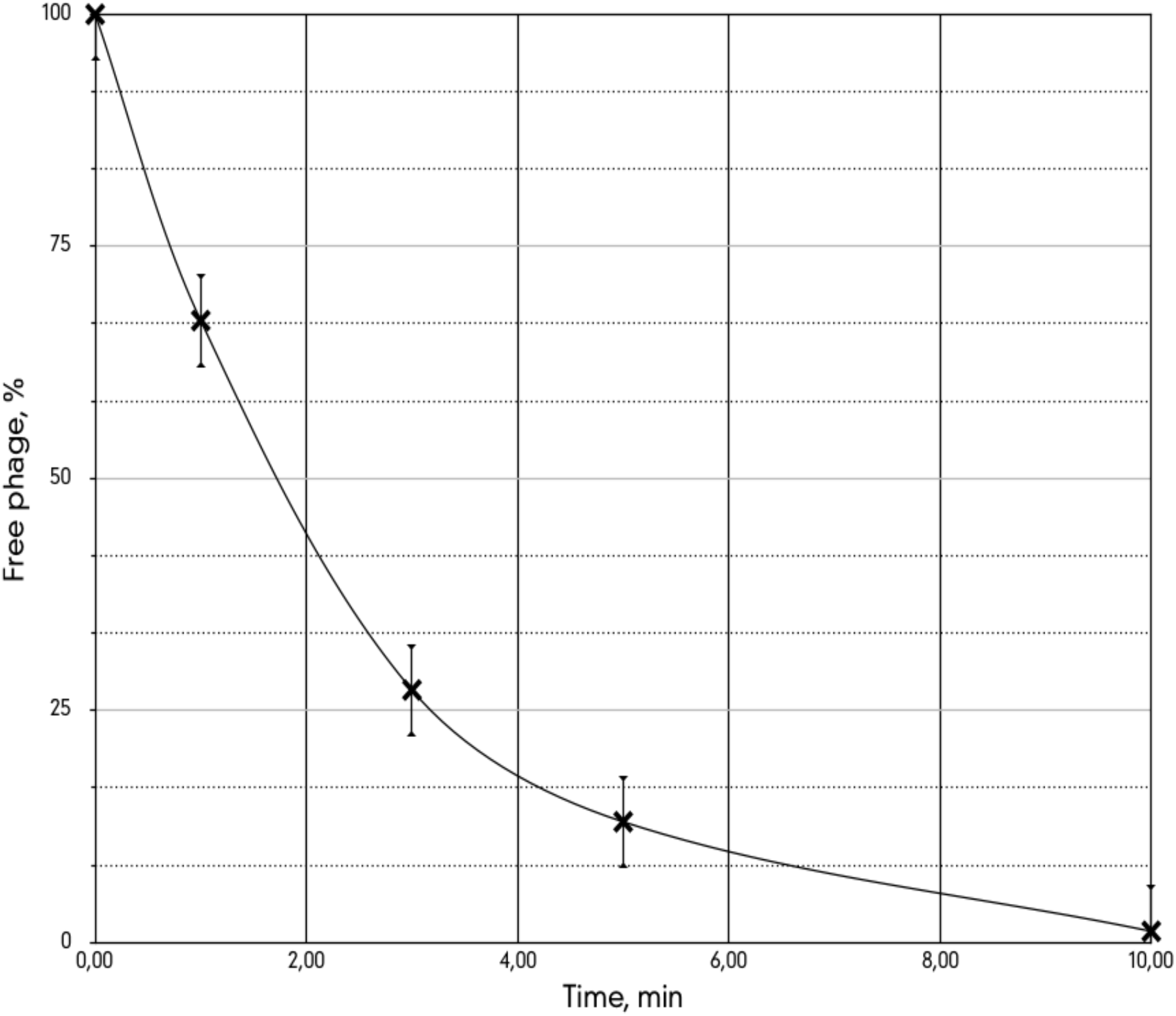
Adsorption curve of bacteriophage Hf4s on *E. coli* 4s cells. The bacterial density was 7×10^7^ ml^−1^, experiments were triplicated.

We searched for possible determinant of the specificity towards the host O antigen in the phage Hf4s genome. In the related bacteriophage P22 the tail spike protein is responsible for the O antigen recognition (49). The same is true for the most of other podoviruses (see (50) for review). The tail spike related gene, g15, was detected in the Hf4s genome (Fig. 2). The C-terminal part of the Hf4s gp15 was related (45% a.a. identity) to the tail spike protein of the T7-related *Escherichia* phage Peacock that was isolated in USA using the same *E. coli* 4s strain as a host. Since *E. coli* 4s was shown to possess O22-like O antigen providing efficient shielding of the intimate outer membrane surface from the interaction with bacteriophage receptor recognition proteins (29, 30, 32) we concluded that the Hf4s tail spike recognizes the host O antigen as the phage primary receptor as similarly to the phage P22 tail spike (51, 52). The HHpred search revealed the structural similarity of the Hf4s tail spike with several phage proteins having the glycosidase activity that is compatible with the polysaccharide receptor recognition.

### 3.5 Lysogen O-seroconversion by the phage Hf4s

The analysis of Hf4s genome revealed the presence of the O antigen seroconversion cluster Gtr consisting of the membrane protein GtrC, glycosyltransferase GtrB and flippase. The Hf4s-encoded GtrC protein was distantly related to *Salmonella* phage Epsilon 34 protein. The glycosyltransferase sequence closely matches several seroconverting *Shigella* phages, for example phage SfX GtrB protein (90% a.a. identity). The flippase also matches *Shigella* phages, as well as the paradigmal serotype converting phage P22 (the best hit is for the phage SfV GtrA with 92% a.a. identity). We compared the adsorption of Hf4s on the parental *E. coli* 4s and on the lysogenic *E. coli* 4s (Hf4s) strain. While 80% of the phage was adsorbed after 10 min of the contact with the host strain mid-log culture (OD_600_ = 0.6), no phage adsorption was detected in the similar conditions with the lysogenic bacteria. This result indicates that the lysogenization by the phage Hf4s leads to the alterations of the O polysaccharide structure and thus to the superinfection exclusion. To further test this hypothesis, we then checked the sensitivity of the lysogen to the bacteriophages known to specifically recognize *E. coli* 4s O antigen (Table 1). The lysogen was sensitive to the bacteriophages G7C and phiKT that not only recognize the O polysaccharide but are strictly dependent on it for the infection (30, 31). At the same time the lysogenic strain was resistant to the T5-like bacteriophage DT57C. This phage is able to infect multiple rough *E. coli* strains *via* the direct recognition of its secondary protein receptor BtuB, however the O antigen of *E. coli* 4s prevents such a contact. For efficient infection of this strain, phage DT57C needs to specifically bind to the O antigen by its lateral tail fiber (LTF) protein LtfA (32). To rule out the possibility of the post-infection exclusion of the DT57C phage by the Hf4s prophage, we selected the lysogenic strain for the resistance to the bacteriophage G7C to produce E. coli (Hf4s) – G7C^R^ strain. It was previously shown (29, 30) that G7C – resistant clones deficient of O antigen synthesis or O antigen O-acetylation. In both cases the non-specific protection of the cell surface gets impaired allowing T5-like bacteriophages to infect such strains independently of the LTF function (29, 32). The *E. coli* 4s (Hf4s)-G7C^R^ lysogens gained the sensitivity to DT57C infection. These clones were also sensitive to the phage FimX, the mutant of DT57C-like phage, devoid of lateral tail fibers. At the same time, the *E. coli* 4s (Hf4s)-G7C^R^ lost the sensitivity to the phage phiKT (Table 1), that is indicative for the complete loss of the O antigen. Therefore, we concluded that the intracellular development of the phage DT57C is not affected by the presence of the Hf4s prophage, however the O antigen of the lysogenic strain is altered and the LTFs of DT57C could no longer recognize it.

We compared the lipopolysaccharide (LPS) profiles of the (Hf4s) lysogen to the parental strain and E. coli 4s GTR^−^ strain, strain using SDS-PAGE electrophoresis (as described in (30)). *E. coli* 4s GTR^−^ strain, lacking the lateral glycosylation had more pronounced downward shift of the LPS bands for about ½ period of LPS stairs-like pattern (30) and Fig. 7), however the lysogen showed no visible bands shift though all the indirect data mentioned above suggested that the O antigen undergoes some structural alterations.

**Figure 7.**
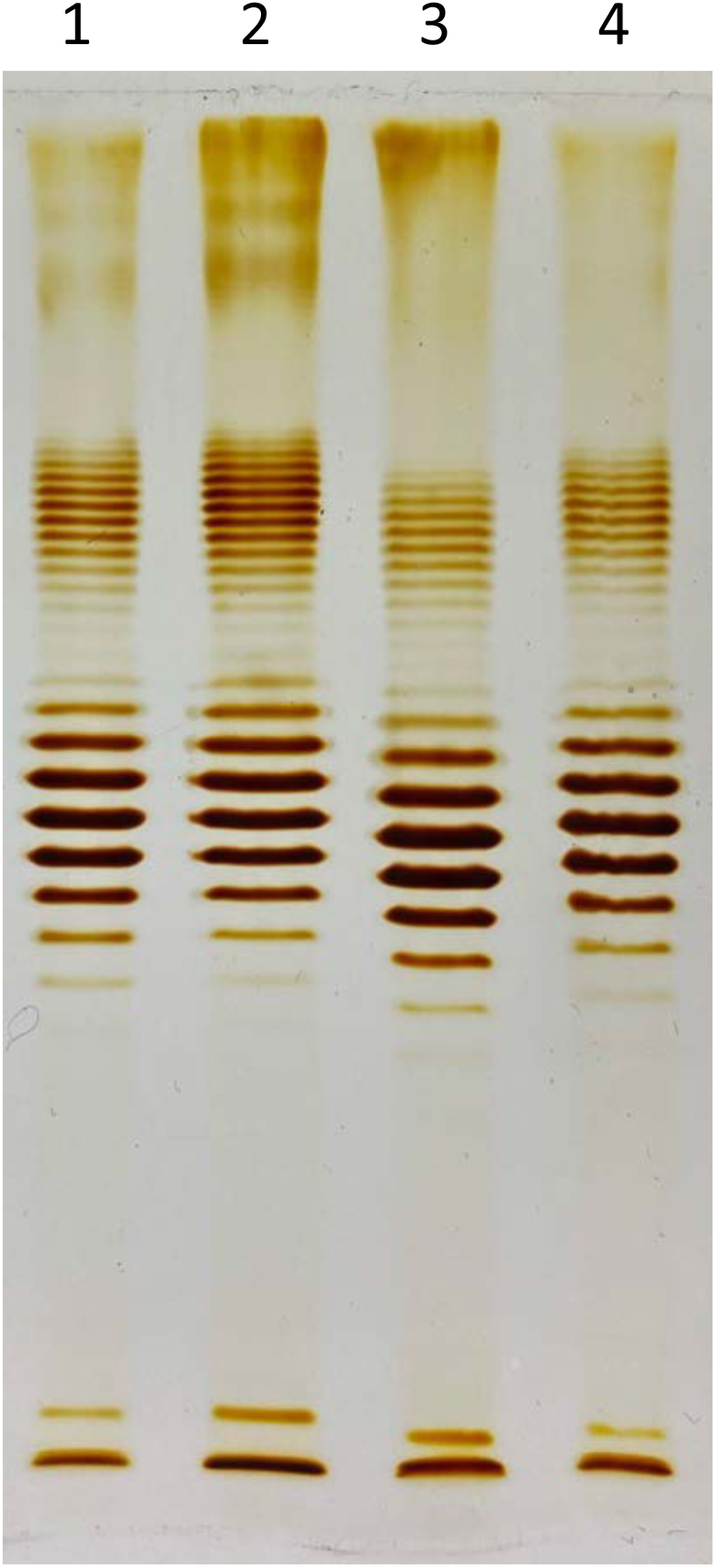
SDS – PAGE electrophoresis of the LPS of *E. coli* 4s (lanes 1 and 4), E. coli 4s (Hf4s) lysogen (lane 2) and *E. coli* 4s GTR mutant, lacking the O-unit lateral glycosylation (lane 3). No difference in band mobility between E. coli 4s and E. coli 4s(Hf4s) can be seen while *E. coli* 4s GTR LPS profile has a marked shift in the bands mobility.

To elucidate precisely the structural changes in the O antigen of the *E. coli* 4s (Hf4s) lysogens, the LPS was extracted from the lysogenic *E. coli* 4s strain and O polysaccharide was isolated by the LPS mild acid degradation. It was studied by NMR spectroscopy essentially as described earlier for the wild-type O polysaccharide of *E. coli* 4s (29). The ^1^H and ^13^C NMR spectra of the O polysaccharide showed signals for anomeric atoms for seven monosaccharide residues (δ_H_ 4.65-5.64, δ_C_ 97.1-103.9) and one O-acetyl group (δ_H_ 2.15, δ_C_ 21.4 and 174.4) demonstrating a mono-O-acetylated heptasaccharide repeating unit. All signals in the NMR spectra were assigned (Table) using two-dimensional NMR spectroscopy, including ^1^H,^1^H COSY, ^1^H,^1^H TOCSY, ^1^H,^1^H ROESY, ^1^H,^13^C HSQC, and ^1^H,^13^C HMBC experiments.

The monosaccharide residues in the O polysaccharide of *E. coli* 4s (Hf4s) were attributed to units **A**-**G**, including three Glc*p* residues (**A**, **F**, and **G**), two residues of Gal*p*NAc (**C** and **E**), and one residue each of Gal*p* (**D**) and ɑ-glucuronic acid (Glc*p*A) (**B**), all being in the pyranose form (Figure). Analysis of intraresidue and interresidue correlations in the two-dimensional ^1^H,^1^H ROESY and ^1^H,^13^C HBMC spectra (data not shown) revealed the same positions and configurations of the glycosidic linkages, sequence of residues **A**-**F**, and O-acetylation pattern (OAc at position 3 of residue **C**; δ_H-3_ 5.13-5.16) in the O polysaccharides of *E. coli* 4s (Hf4s) and *E. coli* 4s (29) (Fig. 8). Based on ^3^*J*_H,H_ coupling constants determined from the ^1^H,^1^H COSY spectrum, the additional monosaccharide residue **G** in the *E. coli* 4s (Hf4s) O polysaccharide was identified as ɑ- Glc*p.*

**Figure 8.**
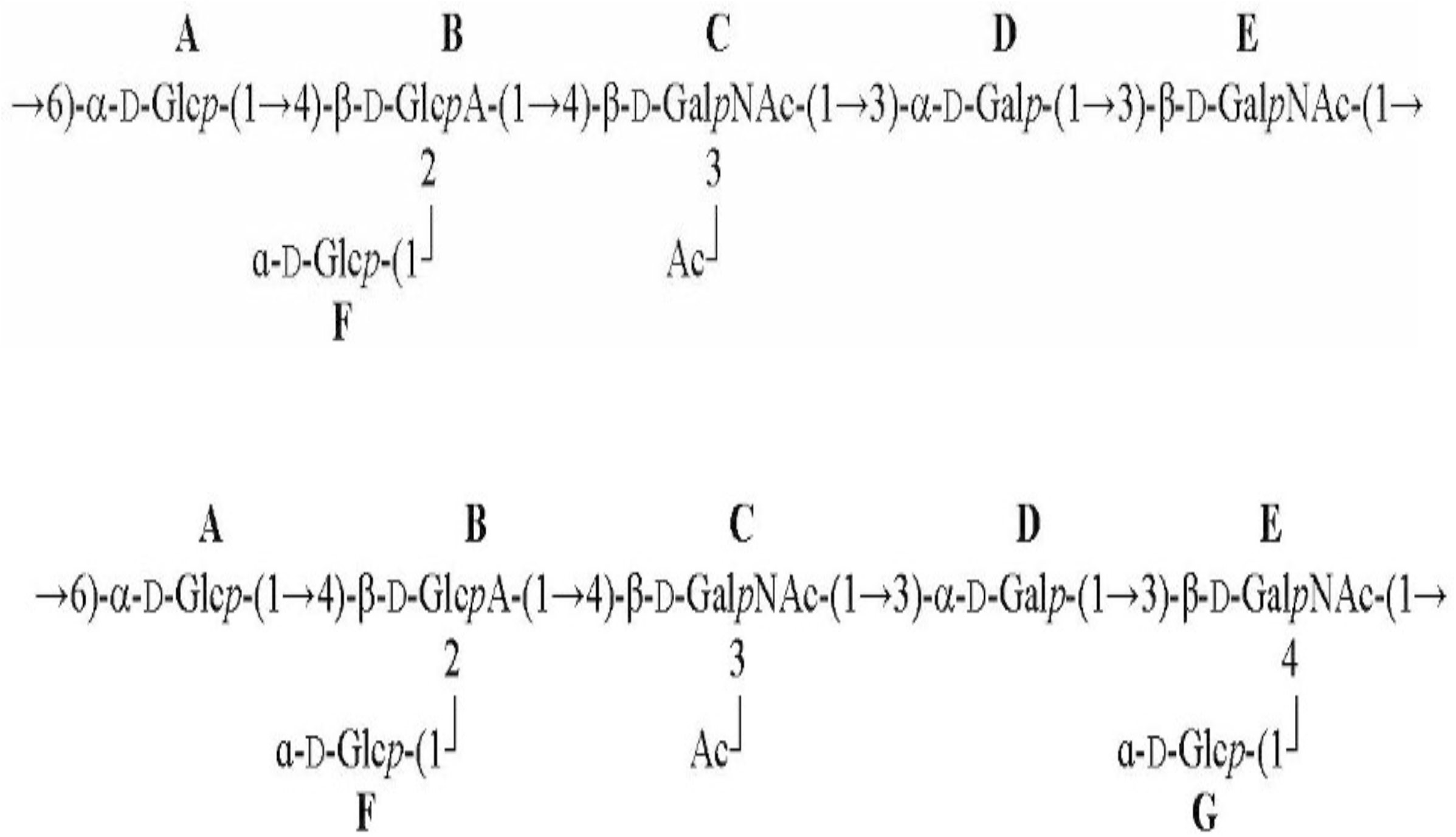
Structure of the O-units of *E. coli* 4s (top) and *E.coli* 4s(Hf4s) lysogen (bottom). Additional lateral glycosylation is present in the O antigen of the lysogenic strain.

A comparison with the NMR data of the O polysaccharides of strains 4s (29) and *E. coli* 4s (Hf4s) (Table S3) showed a close similarity of the ^1^H and ^13^C NMR chemical shifts of five from six common monosaccharide residues (**A**-**D** and **F**). In contrast, the chemical shifts of residue **E** were different, the highest deviation being observed for the C-4 signal (δ_C_ 77.2 in strain *E. coli* 4s (Hf4s) versus δ_C_ 65.3 in strain 4s). Such a significant downfield displacement could be only accounted for by glycosylation of residue **E** at position 4 (53) with residue **G** in the O polysaccharide of strain *E. coli* 4s (Hf4s). The site of attachment of residue **G** was confirmed by a **G** H-1/**E** H-4 cross-peak at δ 5.05/4.31 in the ^1^H,^1^H ROESY spectrum as well as **G** H-1/**E** C-4 and **G** C-1/**E** H-4 correlations at δ 5.05/77.2 and 102.1/ 4.31, respectively, revealed by the ^1^H,^13^C HBMC spectrum.

Monomeric composition, substitution pattern, anomeric absolute and ringsize configurations, sequence of residues and localization of acetyl groups were fed to GODDESS ^13^C, ^1^H, and HSQC NMR simulation service (54), and the comparison of the experimental chemical shifts to the simulated ones confirmed the full structure.

Based on these data, it was concluded that the mutant O polysaccharide of *E. coli* 4s (Hf4s) has the structure shown on the Fig 9. It differs from the wild-type O polysaccharide of *E. coli* 4s only by the presence of the second side-chain glucose residue (**G**).

## 4 Discussion

The intestinal coliphages community of domestic horses, at least in the conditions of the city equestrian center stable (where the animals spend most of their time in the boxes, their diet consisting of oats along with the forages) is heavily dominated by the virulent bacteriophages (11, 24–26, 55). Nevertheless, bacteriophage Hf4s isolated from the horse feces infects an indigenous equine *E. coli* strain and is well adapted to this host since it is not only able to recognize its O antigen as the receptor, but also carries the O-seroconversion module active in *E. coli* 4s (Hf4s) lysogens. This allows us to consider Hf4s as an indigenous component of the equine intestinal microbiome. Up to our knowledge, it is the first report of the isolation of a temperate phage from the horse feces.

Interestingly, the isolation of the bacteriophages recognizing the O antigen of *E.coli* 4s (or structurally similar O antigens of some other *E. coli* strains) from the feces of the horses of the same population (equestrian center Neskuchny sad in Moscow, Russia) was reported for multiple times (10, 29, 32, 56). Some of these phages such as DT57C are restricted by the Hf4s-mediated seroconversion while the others such as G7C or PhiKT are not. It has been shown that the intestinal *E. coli* populations of the horses held at this location were extremely diverse at the strain level with up to 1000 of distinct *E. coli* strain present in the same sample of the feces. It was also suggested that the phage infection pressure might contribute to creation and maintenance of such a high intraspecies diversity (11, 57). This allows us to speculate that the acquisition of Hf4s prophage may have ecological significance as an adaptation of *E. coli* to co-existence with virulent coliphages (11) and, on the other hand, it may influence the ecology of some virulent phages, especially of those featuring narrow host ranges been dependent on this particular O antigen type for infection (30).

Under the laboratory conditions, the bacteriophage Hf4s produces rather large plaques with marked concentric rings pattern that is more pronounced than it was observed in lambda phage. The morphology of Hf4s plaques appears to be closer to lambda CIII mutants (58). However, in our case the wild type phage is used. It is possible that the lysogens surface modification may contribute to this phenomenon, though more experiments are needed to elucidate the underlying mechanism. Due to well visible ring pattern phage Hf4s – *E. coli* 4s system may turn a useful model object for deciphering of the mechanisms of the morphogenesis of the complex spatial patterns emerging due to the bacteriophage infection, which may have certain significance not only in the laboratory but also in some natural habitats.

Summarizing all the data, we conclude that Hf4s is a temperate bacteriophage infecting indigenous equine intestinal *E. coli* isolate that based on its genomic features can be classified within the *Lederbergvirus* genus. Phage Hf4s carries a functional O-antigen seroconversion cluster that alters the recognition of the lysogen’s cell surface by some coliphages and may thus contribute to the host fitness in the horse intestinal ecosystem.

## Author Contribution

Conceptualization, Alla Golomidova; Data curation, Alexandr Efimov, Ilya Belalov; Funding acquisition, Andrey Letarov; Investigation, Alla Golomidova; Alexandr Efimov, Evelina Zdorovenko, Yuri Knirel, Maria Letarova, Methodology, Eugene Kulikov; Project administration, Andrey Letarov; Supervision, Eugene Kulikov; Validation, Alexandr Efimov, Ilya Belalov; Writing – original draft, Andrey Letarov; Writing – review & editing, Eugene Kulikov. All the authors read and edited the manuscript.

## Acknowledgements

The authors are grateful to Vladislav Babenko for the help with phage genomic sequencing, to Natalia Pushkina for phage adsorption experiments and to Roman Belousoff for taking part in material collection.

---

**Figure S1.**
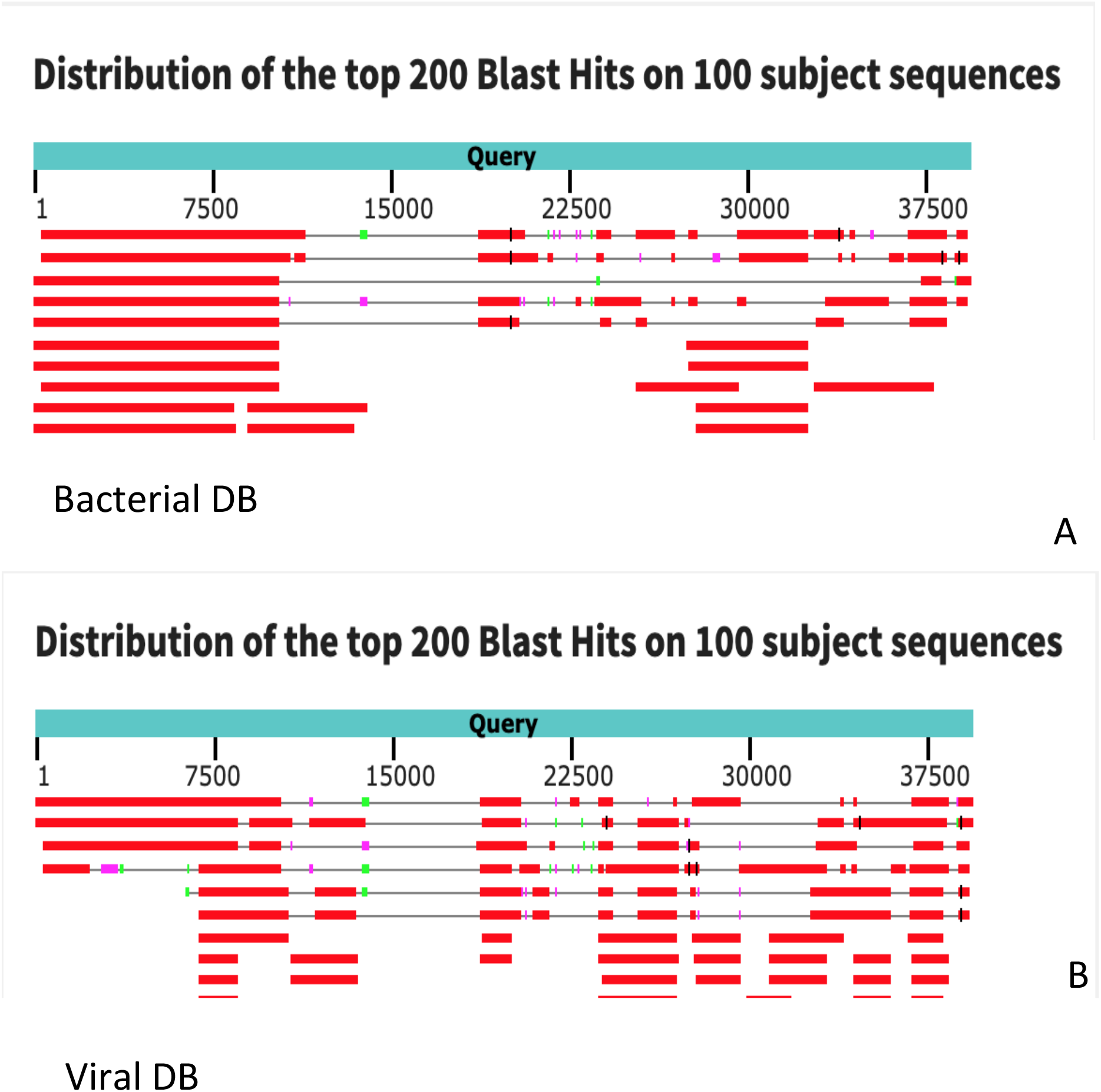
The results of blastN analysis of the phage Hf4s whole genome sequence. A – The output of the nBlast run against the viral database. Ten best hits are shown. B – Prophages, related to Hf4s.

## References

1. Letarov A, Kulikov E. 2009. The bacteriophages in human- and animal body-associated microbial communities. J Appl Microbiol 107:1–13.

2. Moreno-Gallego JL, Chou SP, Di Rienzi SC, Goodrich JK, Spector TD, Bell JT, Youngblut ND, Hewson I, Reyes A, Ley RE. 2019. Virome Diversity Correlates with Intestinal Microbiome Diversity in Adult Monozygotic Twins. Cell Host Microbe 25:261–272 e5.

3. Seo SU, Kweon MN. 2019. Virome-host interactions in intestinal health and disease. Curr Opin Virol 37:63–71.

4. Shkoporov AN, Hill C. 2019. Bacteriophages of the Human Gut: The “Known Unknown” of the Microbiome. Cell Host Microbe 25:195–209.

5. Sausset R, Petit MA, Gaboriau-Routhiau V, De Paepe M. 2020. New insights into intestinal phages. Mucosal Immunol 13:205–215.

6. Babenko VV, Millard A, Kulikov EE, Spasskaya NN, Letarova MA, Konanov DN, Belalov IS, Letarov AV. 2020. The ecogenomics of dsDNA bacteriophages in feces of stabled and feral horses. Comput Struct Biotechnol J 18:3457–3467.

7. Guerin E, Shkoporov AN, Stockdale SR, Comas JC, Khokhlova EV, Clooney AG, Daly KM, Draper LA, Stephens N, Scholz D, Ross RP, Hill C. 2021. Isolation and characterisation of PhicrAss002, a crAss-like phage from the human gut that infects Bacteroides xylanisolvens. Microbiome 9:89.

8. Cervantes-Echeverria M, Equihua-Medina E, Cornejo-Granados F, Hernandez-Reyna A, Sanchez F, Lopez-Contreras BE, Canizales-Quinteros S, Ochoa-Leyva A. 2018. Whole-genome of Mexican-crAssphage isolated from the human gut microbiome. BMC Res Notes 11:902.

9. Mathieu A, Dion M, Deng L, Tremblay D, Moncaut E, Shah SA, Stokholm J, Krogfelt KA, Schjorring S, Bisgaard H, Nielsen DS, Moineau S, Petit MA. 2020. Virulent coliphages in 1-year-old children fecal samples are fewer, but more infectious than temperate coliphages. Nat Commun 11:378.

10. Babenko V, Golomidova A, Ivanov P, Letarova M, Kulikov E, Manolov A, Prokhorov N, Kostrukova E, Matyushkina D, Prilipov A. 2019. Phages associated with horses provide new insights into the dominance of lateral gene transfer in virulent bacteriophages evolution in natural systems. Biorxiv doi:10.1101/542787:542787.

11. Golomidova A, Kulikov E, Isaeva A, Manykin A, Letarov A. 2007. The diversity of coliphages and coliforms in horse feces reveals a complex pattern of ecological interactions. Appl Environ Microbiol 73:5975–81.

12. Howard-Varona C, Hargreaves KR, Abedon ST, Sullivan MB. 2017. Lysogeny in nature: mechanisms, impact and ecology of temperate phages. ISME J 11:1511–1520.

13. Baxter JC, Waples WG, Funnell BE. 2020. Nonspecific DNA binding by P1 ParA determines the distribution of plasmid partition and repressor activities. J Biol Chem 295:17298–17309.

14. Hammerl JA, Jackel C, Funk E, Pinnau S, Mache C, Hertwig S. 2016. The diverse genetic switch of enterobacterial and marine telomere phages. Bacteriophage 6:e1148805.

15. Samson JE, Magadan AH, Sabri M, Moineau S. 2013. Revenge of the phages: defeating bacterial defences. Nat Rev Microbiol 11:675–87.

16. Vander Byl C, Kropinski AM. 2000. Sequence of the genome of Salmonella bacteriophage P22. J Bacteriol 182:6472–81.

17. Taylor VL, Fitzpatrick AD, Islam Z, Maxwell KL. 2019. The Diverse Impacts of Phage Morons on Bacterial Fitness and Virulence. Adv Virus Res 103:1–31.

18. Feiner R, Argov T, Rabinovich L, Sigal N, Borovok I, Herskovits AA. 2015. A new perspective on lysogeny: prophages as active regulatory switches of bacteria. Nat Rev Microbiol 13:641–50.

19. Holt GS, Lodge JK, McCarthy AJ, Graham AK, Young G, Bridge SH, Brown AK, Veses-Garcia M, Lanyon CV, Sails A, Allison HE, Smith DL. 2017. Shigatoxin encoding Bacteriophage varphi24B modulates bacterial metabolism to raise antimicrobial tolerance. Sci Rep 7:40424.

20. Bobay LM, Touchon M, Rocha EP. 2014. Pervasive domestication of defective prophages by bacteria. Proc Natl Acad Sci U S A 111:12127–32.

21. Reyes A, Haynes M, Hanson N, Angly FE, Heath AC, Rohwer F, Gordon JI. 2010. Viruses in the faecal microbiota of monozygotic twins and their mothers. Nature 466:334–8.

22. Furuse K, Osawa S, Kawashiro J, Tanaka R, Ozawa A, Sawamura S, Yanagawa Y, Nagao T, Watanabe I. 1983. Bacteriophage distribution in human faeces: continuous survey of healthy subjects and patients with internal and leukaemic diseases. J Gen Virol 64 (Pt 9):2039–43.

23. Chibani-Chennoufi S, Sidoti J, Bruttin A, Dillmann ML, Kutter E, Qadri F, Sarker SA, Brussow H. 2004. Isolation of Escherichia coli bacteriophages from the stool of pediatric diarrhea patients in Bangladesh. J Bacteriol 186:8287–94.

24. Kulikov E, Kropinski AM, Golomidova A, Lingohr E, Govorun V, Serebryakova M, Prokhorov N, Letarova M, Manykin A, Strotskaya A, Letarov A. 2012. Isolation and characterization of a novel indigenous intestinal N4-related coliphage vB_EcoP_G7C. Virology 426:93–9.

25. Kulikov EE, Golomidova AK, Letarova MA, Kostryukova ES, Zelenin AS, Prokhorov NS, Letarov AV. 2014. Genomic sequencing and biological characteristics of a novel Escherichia coli bacteriophage 9g, a putative representative of a new Siphoviridae genus. Viruses 6:5077–92.

26. Golomidova AK, Kulikov EE, Prokhorov NS, Guerrero-Ferreira RC, Ksenzenko VN, Tarasyan KK, Letarov AV. 2015. Complete genome sequences of T5-related Escherichia coli bacteriophages DT57C and DT571/2 isolated from horse feces. Arch Virol 160:3133–7.

27. Golomidova AK, Kulikov EE, Babenko VV, Kostryukova ES, Letarov AV. 2018. Complete Genome Sequence of Bacteriophage St11Ph5, Which Infects Uropathogenic Escherichia coli Strain up11. Genome Announc 6.

28. Zdorovenko EL, Wang Y, Shashkov AS, Chen T, Ovchinnikova OG, Liu B, Golomidova AK, Babenko VV, Letarov AV, Knirel YA. 2018. O-Antigens of Escherichia coli Strains O81 and HS3-104 Are Structurally and Genetically Related, Except O-Antigen Glucosylation in E. coli HS3-104. Biochemistry (Mosc) 83:534–541.

29. Knirel YA, Prokhorov NS, Shashkov AS, Ovchinnikova OG, Zdorovenko EL, Liu B, Kostryukova ES, Larin AK, Golomidova AK, Letarov AV. 2015. Variations in O-antigen biosynthesis and O-acetylation associated with altered phage sensitivity in Escherichia coli 4s. J Bacteriol 197:905–12.

30. Kulikov EE, Golomidova AK, Prokhorov NS, Ivanov PA, Letarov AV. 2019. High-throughput LPS profiling as a tool for revealing of bacteriophage infection strategies. Sci Rep 9:2958.

31. Prokhorov NS, Riccio C, Zdorovenko EL, Shneider MM, Browning C, Knirel YA, Leiman PG, Letarov AV. 2017. Function of bacteriophage G7C esterase tailspike in host cell adsorption. Mol Microbiol 105:385–398.

32. Golomidova AK, Kulikov EE, Prokhorov NS, Guerrero-Ferreira RC, Knirel YA, Kostryukova ES, Tarasyan KK, Letarov AV. 2016. Branched Lateral Tail Fiber Organization in T5-Like Bacteriophages DT57C and DT571/2 is Revealed by Genetic and Functional Analysis. Viruses 8.

33. Bankevich A, Nurk S, Antipov D, Gurevich AA, Dvorkin M, Kulikov AS, Lesin VM, Nikolenko SI, Pham S, Prjibelski AD, Pyshkin AV, Sirotkin AV, Vyahhi N, Tesler G, Alekseyev MA, Pevzner PA. 2012. SPAdes: a new genome assembly algorithm and its applications to single-cell sequencing. J Comput Biol 19:455–77.

34. Camacho C, Coulouris G, Avagyan V, Ma N, Papadopoulos J, Bealer K, Madden TL. 2009. BLAST+: architecture and applications. BMC Bioinformatics 10:421.

35. Starikova EV, Tikhonova PO, Prianichnikov NA, Rands CM, Zdobnov EM, Ilina EN, Govorun VM. 2020. Phigaro: high-throughput prophage sequence annotation. Bioinformatics 36:3882–3884.

36. Sullivan MJ, Petty NK, Beatson SA. 2011. Easyfig: a genome comparison visualizer. Bioinformatics 27:1009–10.

37. Darling AE, Mau B, Perna NT. 2010. progressiveMauve: multiple genome alignment with gene gain, loss and rearrangement. PLoS One 5:e11147.

38. Meier-Kolthoff JP, Auch AF, Klenk HP, Goker M. 2013. Genome sequence-based species delimitation with confidence intervals and improved distance functions. BMC Bioinformatics 14:60.

39. Meier-Kolthoff JP, Goker M. 2017. VICTOR: genome-based phylogeny and classification of prokaryotic viruses. Bioinformatics 33:3396–3404.

40. Lefort V, Desper R, Gascuel O. 2015. FastME 2.0: A Comprehensive, Accurate, and Fast Distance-Based Phylogeny Inference Program. Mol Biol Evol 32:2798–800.

41. Farris JS. 1972. Estimating phylogenetic trees from distance matrices. The American Naturalist 106:645–668.

42. Rambaut A. 2009. FigTree. Tree figure drawing tool. http://tree.bio.ed.ac.uk/software/figtree/. Accessed May 8.

43. Goker M, Garcia-Blazquez G, Voglmayr H, Telleria MT, Martin MP. 2009. Molecular taxonomy of phytopathogenic fungi: a case study in Peronospora. PLoS One 4:e6319.

44. Meier-Kolthoff JP, Hahnke RL, Petersen J, Scheuner C, Michael V, Fiebig A, Rohde C, Rohde M, Fartmann B, Goodwin LA, Chertkov O, Reddy T, Pati A, Ivanova NN, Markowitz V, Kyrpides NC, Woyke T, Goker M, Klenk HP. 2014. Complete genome sequence of DSM 30083(T), the type strain (U5/41(T)) of Escherichia coli, and a proposal for delineating subspecies in microbial taxonomy. Stand Genomic Sci 9:2.

45. Sauvageau D, Cooper D. 2010. Two-stage, self-cycling process for the production of bacteriophages. Microbial Cell Factories 9:81.

46. L’vov VL, Dashunin VM, Ramos EL, Shashkov AS, Dmitriev BA, Kochetkov NK. 1983. Somatic antigens of Shigella: the structure of the polysaccharide chain of Shigella boydii type 2 lipopolysaccharide. Carbohydrate research 124:141–149.

47. de Siqueira RS, Dodd CE, Rees CE. 2006. Evaluation of the natural virucidal activity of teas for use in the phage amplification assay. Int J Food Microbiol 111:259–62.

48. Juhala RJ, Ford ME, Duda RL, Youlton A, Hatfull GF, Hendrix RW. 2000. Genomic sequences of bacteriophages HK97 and HK022: pervasive genetic mosaicism in the lambdoid bacteriophages. J Mol Biol 299:27–51.

49. Wang G, Jin S, Ling X, Li Y, Hu Y, Zhang Y, Huang Y, Chen T, Lin J, Ning Z, Meng Y, Li X. 2019. Proteomic Profiling of LPS-Induced Macrophage-Derived Exosomes Indicates Their Involvement in Acute Liver Injury. Proteomics 19:e1800274.

50. Letarov AV, Kulikov EE. 2017. Adsorption of Bacteriophages on Bacterial Cells. Biochemistry (Mosc) 82:1632–1658.

51. Walter M, Fiedler C, Grassl R, Biebl M, Rachel R, Hermo-Parrado XL, Llamas-Saiz AL, Seckler R, Miller S, van Raaij MJ. 2008. Structure of the receptor-binding protein of bacteriophage det7: a podoviral tail spike in a myovirus. Journal of virology 82:2265–73.

52. Andres D, Baxa U, Hanke C, Seckler R, Barbirz S. 2010. Carbohydrate binding of Salmonella phage P22 tailspike protein and its role during host cell infection. Biochemical Society Transactions 38:1386–1389.

53. Shashkov AS, Lipkind GM, Knirel YA, Kochetkov NK. 1988. Stereochemical factors determining the effects of glycosylation on the ^13^C chemical shifts in carbohydrates. Magnetic resonance in chemistry 26:735–747.

54. Kapaev RR, Toukach PV. 2016. Simulation of 2D NMR spectra of carbohydrates using GODESS software. Journal of Chemical Information and Modeling 56:1100–1104.

55. Golomidova AK, Kulikov EE, Babenko VV, Ivanov PA, Prokhorov NS, Letarov AV. 2019. Escherichia coli bacteriophage Gostya9, representing a new species within the genus T5virus. Archives of virology 164:879–884.

56. Golomidova AK, Kulikov EE, Kudryavtseva AV, Letarov AV. 2018. Complete Genome Sequence of Escherichia coli Bacteriophage PGT2. Genome Announc 6.

57. Isaeva AS, Kulikov EE, Tarasyan KK, Letarov AV. 2010. A Novel High-Resolving Method for Genomic PCR-Fingerprinting of Enterobacteria. Acta Naturae 2:82–8.

58. Mitarai N, Brown S, Sneppen K. 2016. Population Dynamics of Phage and Bacteria in Spatially Structured Habitats Using Phage lambda and Escherichia coli. J Bacteriol 198:1783–93.

